# Categorical working memory codes in human visual cortex

**DOI:** 10.1101/2021.12.16.472288

**Authors:** Chang Yan, Thomas B. Christophel, Carsten Allefeld, John-Dylan Haynes

## Abstract

Working memory contents are represented in neural activity patterns across multiple regions of the cortical hierarchy. It has remained unclear to which degree this reflects a specialization for different levels of abstraction. Here, we demonstrate that for color stimuli categorical codes are already present at the level of extrastriate visual cortex (V4 and VO1). Interestingly, this categorical coding was observed during working memory, but not during perception.

---

The human mind has the ability to temporally store sensory information to guide decision making and behavior^1^. Information about memorized sensory contents has been found in activity patterns across numerous cortical regions^2^. Importantly, the same memorized content can be represented in more than one region^3–9^, even at the same time^10^. While these multiple representations might be simply redundant, it has been suggested that there could be a division of labor such that early sensory regions encode low-level sensory details whereas more higher-level regions represent increasingly abstract and categorical properties of memorized stimuli^2,11^. To date, it has remained unclear at which stage the categorical nature of memory representations begins to emerge.

Color stimuli have long been used to study the capacity and precision of visual working memory^12–16^. Importantly, color stimuli exhibit both continuous and categorical properties. For example, colors are perceived as a continuum but they are also readily grouped into basic color categories^17,18^, even when patients are incapable of naming them^19^.

Recent behavioral work suggested that performance during continuous color recall could be explained by a dual content model that combines categorical and continuous (non-categorical) components^20,21^. Such a dual content model reliably predicted color reproduction that were consistently biased away from the memorized color.

Here we used fMRI and multivariate encoding models to assess whether brain representations of memorized colors in different visual brain areas are better explained by more continuous or more categorical codes. To this end, we scanned 10 healthy participants in multiple MRI sessions (4 each). They performed a conventional color working memory task requiring subjects to recall a remembered color as accurately as possible and indicate their choice with a continuous color wheel. Subjects either recalled the colors immediately (undelayed ’perceptual’ task, see **Figure 1b**) or after a delay (delayed ’memory’ task, see **Figure 1a**). To closely capture the neural activity patterns encoding colors of different hues, we sampled colors evenly from a calibrated color space.

**Figure 1.**
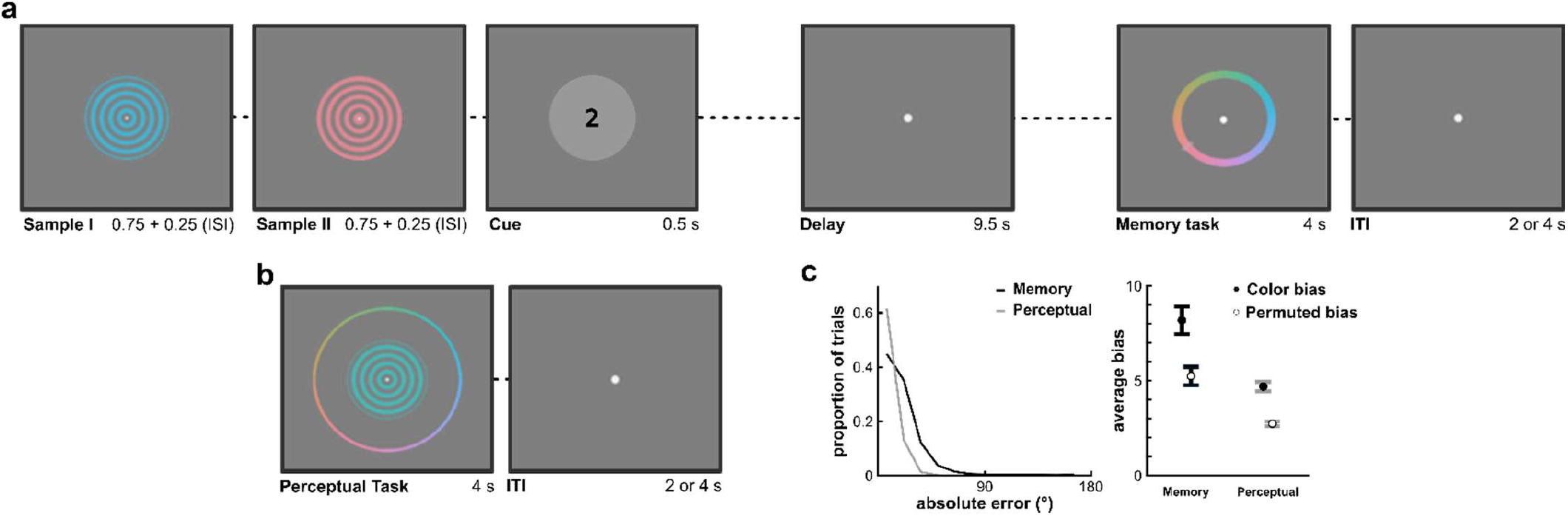
Experimental design and stimuli. **(a)** In the working memory task, subjects were presented with two color samples followed by a retro-cue (‘1’ or ‘2’). The retro-cue indicated which of the two items had to be recalled by clicking on the respective color on a color wheel after the delay (memory task). **(b)** In the perceptual task, the sample and the color wheel were shown at the same time minimizing mnemonic demands. **(c)** Subjects made larger errors in the memory task (absolute error in degrees of the color-wheel, collapsed across subjects) and showed more consistent biases in color reports for individual hues (bias defined as absolute error for each color, averaged and compared to biases for permuted color labels, see Methods for details).

## RESULTS

Subjects made larger errors in the memory task (mean absolute error = 17.47° ±1.52° SEM in degrees of the color-wheel, see **Figure 1c**) than in the perceptual task (mean absolute error = 9.8°±0.39° SEM, Wilcoxon signed rank test, p = 0.002), and showed larger categorical biases independently of the overall effect of the errors (see **Figure 1c**, Wilcoxon signed rank test, p = 0.02).

In a behavioral session after the fMRI experiments, we used two separate category-based behavioral tasks to obtain a categorical model of color representation. Following prior behavioral work^20^, subjects performed two tasks: In the color identification task they were given a color name and asked to identify that color on a continuous color wheel (see **Figure 2a**). In the color naming task subjects assigned a color name to a continuous color (see **Figure 2b**). We used 7 color names for these tasks but only six were consistently used by the participants in the naming task. As in prior work^20^, data for the seventh, unused color name (‘red’) was discarded. The resulting behavioral data allow to assess two properties of color representation: The boundaries between color names and the prototypical exemplars for the six color categories most commonly used (see **Figure 2d** and Methods for details). We then averaged the underlying naming and identification distributions to form a simplified categorical encoding model (see **Figure 2c**). Such a model has been used in the past to predict behavioral bias in color recall^20^, but serves here as an approximation of the tuning of category-selective neurons for these color categories responding strongest to the prototypical color and showing a sharp decline towards the boundaries. For comparison we used a standard cosine-shaped continuous sensory (non-categorical) encoding model (see **Figure 2e**), versions of which have been used in previous investigations of working memory coding and precision^6,9,22,23^. This sensory model did not consider information about the delineations of common color categories.

**Figure 2.**
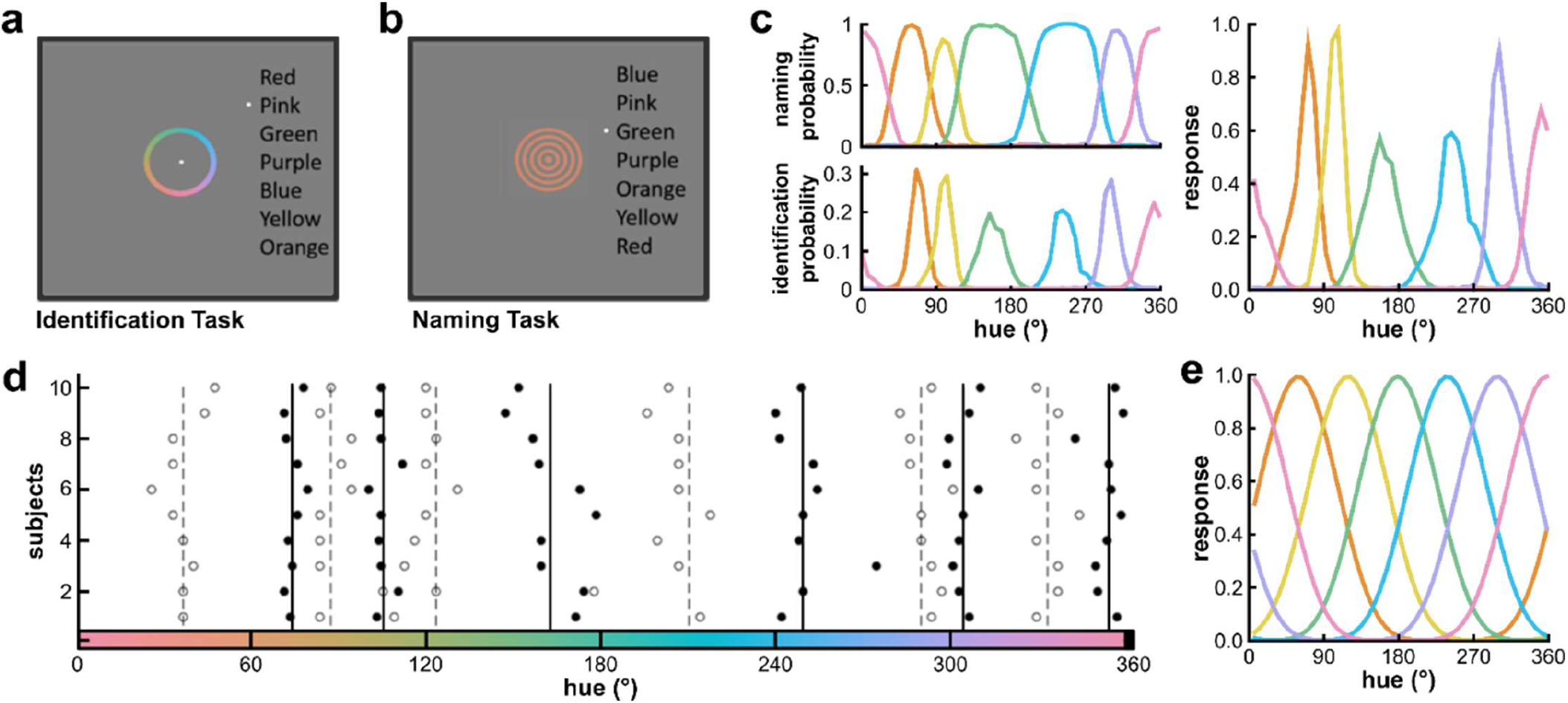
Behavioral categorization tasks and resulting encoding models. We used two category-based tasks performed in a separate session post-MRI where subjects identified prototypical colors on a color wheel **(a)** and named presented colors **(b)**. **(c)** We combined naming and identification frequencies for individual hues to create a tuning model for voxels selective to six color categories **(as in** ^20^). **(d)** This model allows to estimate the boundaries between color categories (open circles, dashed lines) and the prototypical exemplars for each color category (closed circles, solid lines). **(e)** For comparison, we used a standard uniform model typically thought to capture the sampling of sensory units by an fMRI voxel.

We then tested which of these two models best explained mnemonic representations of color in the brain. Using probabilistic anatomical regions of interest^24^ (see methods for details), we focused our analyses on regions of the visual cortex known to have representations of color (**Figure 3a**; V1, V4 and VO1, ^23^). Notably, we used a cross-validated form of multivariate analysis of variance (cvMANOVA^25^, see methods for details) to assess which of the two models better explained the underlying data instead of inverting the model to reconstruct the memorized or the reported color (see ^26^). This is important as we aimed at identifying intermediary representations that not necessarily match either the encoded or the reported color.

**Figure 3.**
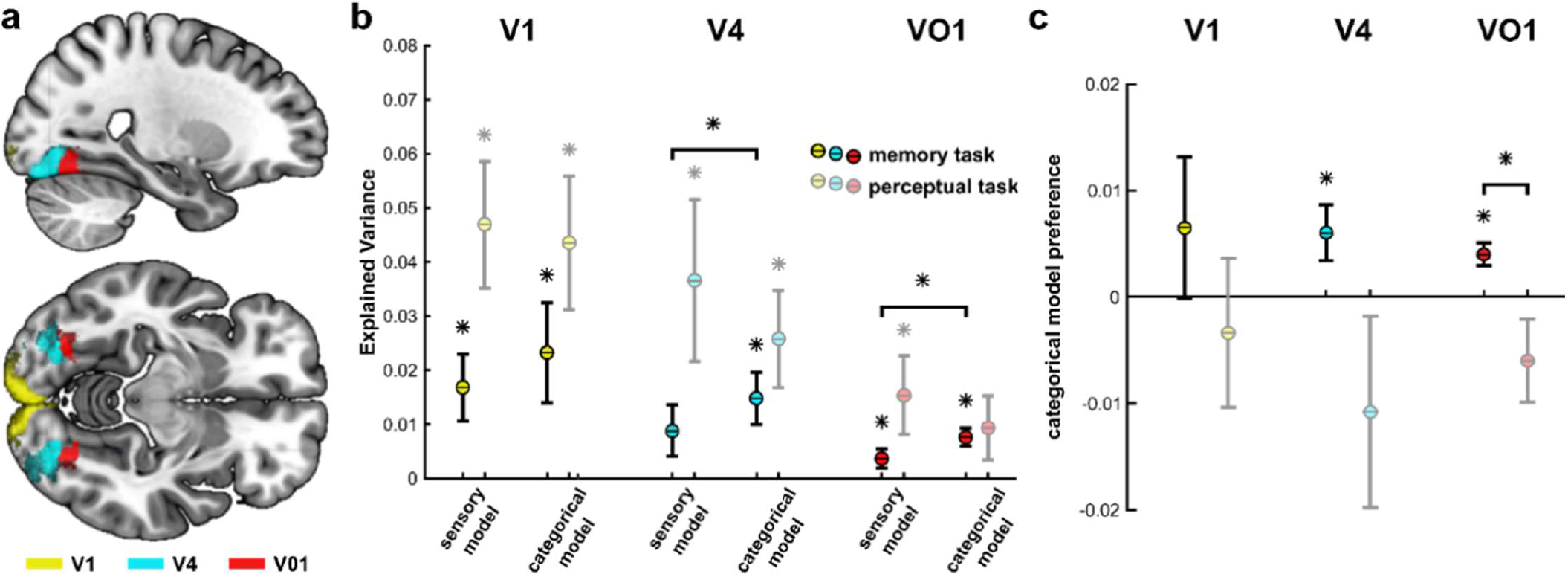
Decoding color-specific information in visual cortex. **(a)** We focused our analyses on anatomically defined regions of interest of V1, V4 and V01. **(b)** Explained variance in fMRI voxel activity patterns for different encoding models (sensory and categorical) during the memory and the immediate task. Regions are color coded; asterisks indicate significance (bootstrapped confidence intervals with p < 0.05; multi-comparison corrected); error bars indicate SEM. **(c)** Categorical model preference in the three regions during the two tasks. Please note that categorical model preferences in V4 are significantly larger during memory as compared to perception when multiple comparisons corrections are omitted (p < 0.05, CI^95^ = [−0.0377, −0.0011]).

Using the sensory model, we found robust representations of the memorized color in V1 and V01 (**Figure 3b**; 95% bootstrapped confidence interval of the explained variance corrected for multiple comparisons; CI^95^ = [0.0017, 0.0292] in V1, CI^95^ = [−0.0038, 0.0181] in V4, CI^95^ = [0.0003, 0.0082] in VO1). When using the categorical model, all three ROIs showed statistically significant information about the target color (CI^95^ = [0.0043, 0.0469] in V1, CI^95^ = [0.0014, 0.0243] in V4, CI^95^ = [0.0041, 0.0124] in VO1).

Notably, the two models are non-orthogonal with respect to each other, meaning that they can be expected to capture shared variance. Hence, we statistically compared which of the two models explained more variance in the underlying data. We found that the categorical color model explained more variance in V4 and V01 than the sensory model that was not informed by the delineations of common color categories (**Error! Reference source not found.c**; 95% bootstrapped confidence interval corrected for multiple comparisons of the categorical model preference; CI^95^ = [-0.009, 0.0212] in V1, CI^95^ = [0.0017, 0.0152] in V4, CI^95^ = [0.0012, 0.0065] in VO1). This suggests that during working memory V4 and V01 encode memoranda in a categorical rather than a sensory format.

Then, we asked how color was represented when subjects did not have to retain the color samples in memory but could immediately report the presented color. For data from this perceptual task, the sensory model explained significant levels of variance in all three regions (**Figure 3b;** CI^95^ = [0.0198, 0.0724] in V1, CI^95^ = [0.0085, 0.0781] in V4, CI^95^ = [0.0013, 0.034] in VO1). In contrast, using the categorical encoding model only V1 and V4 but not VO1 explained above-chance variance (CI^95^ = [0.0202, 0.0783] in V1, CI^95^ = [0.0082, 0.0495] in V4, CI^95^ = [−0.0055, 0.0222] in VO1). Critically, for the perceptual task we found no significant differences in the variance explained by the two models in any region (CI^95^ = [−0.0174, 0.0149] in V1, CI^95^ = [-0.0384, 0.0049] in V4, CI^95^ = [−0.0144, 0.0035] in VO1). Comparing these differences across the two versions of the task (delayed and undelayed) resulted in a significant interaction effect in V01 (**Error! Reference source not found.c**; 95% bootstrapped confidence interval corrected for multiple comparisons of the difference in categorical model preference; CI^95^ = [−0.0238, 0.0118] in V1, CI^95^ = [−0.0429, 0.0016] in V4, CI^95^ = [−0.0177, −0.0023] in VO1) indicating that neural representations in the delayed task were substantially more categorical than in the undelayed task in VO1.

These results suggest that representations of memorized colors are retained by categorical representations already inside visual cortex. This demonstrates that already in visual cortex, different regions retain representations in a differential neural code, and suggests that the memory storage of an individual item does rely on its representation in multiple concurrent coding schemes. Importantly, this preference for categorical tuning is only present during working memory and is absent during an immediate perceptual task. Thus, initially more uniform representations might morph to become more categorical during memory. This could have several potential reasons, such as feedback from higher regions. This finding might serve as a direct explanation for the differential amount of biasing of behavioral responses in immediate and delayed recall in this study and in prior work^20,21^.

In V1, we found that neither model outperformed the other which is consistent with prior work^23^ showing that V1 seems to harbor a fundamentally different discontinuous code (with respect to hue) based on opponent colors which is not reflected by either model. Further research is needed to investigate the vast space of potential color encoding models to give more insight into the properties and determinants of neural coding during working memory and perception.

## Materials and Methods

### Participants

Ten right-handed healthy German native speakers (aged 18-35 years; mean age: 27, SEM ± 1.13; 9 female) with normal or corrected-to-normal vision and no color blindness participated in the study. The sample size and the number of repetitions per task was chosen based on previous studies using similar analyses techniques to study perceptual color representations as well as working memory representations^23,26^, and was considerably increased. We decided to recruit a small subject number with multiple sessions per subject, instead of a large number of subjects^27^. This study was granted ethical approval by the local ethics committee and all subjects gave informed consent.

### Experimental Design

Each subject completed five sessions of experiments, including three 2-h fMRI sessions with 16 runs (50 trials/run) for a delayed estimation task (‘memory task’), one 2-h fMRI session with 14 to 16 runs (50 trials/run) for an undelayed estimation task (‘perceptual task’) both using color as stimulus material (see **Figure 1ab**). The third and last session was one 90-min behavioral session for the color categorization tasks (see **Figure 2ab**). These five sessions were conducted on different days, but within the same month. After the last fMRI session, participants also completed a 2-page questionnaire regarding their strategies for completing the working memory task. All experimental tasks were coded using PsychToolbox-3 (http://psychtoolbox.org/) and MATLAB 2014b (MathWorks, Natick, MA).

### Memory Task

In the delayed estimation task (‘memory task’), subjects memorized a sample hue during a delay period and then reported the memorized color on a randomly rotated color wheel. A trial started with the sequential presentation of two color samples in the middle of the screen, followed by a retro-cue^28^ (either ‘1’ or ‘2’) at the center of a light grey circle (see **Figure 1a**). The sample stimuli were concentric sinusoidal gratings within a circular aperture changing from the central gray point to the sample color, which drifted at a constant speed in a random direction: either inward or outward^23^. A retro-cue informed subjects which of the two sample stimuli should be memorized for the rest of the trial (‘1’ or ‘2’). The retro-cue was followed by the presentation of a blank screen (with only the fixation point) for 9.5 seconds, resulting in an overall delay of 10 s for memorization of the cued stimulus. Then a color wheel included all 50 color samples was presented in the center of the screen. Subjects were asked to indicate on the color wheel which sample sampled they had memorized within 4 s. For this, they scrolled with an MRI compatible trackball from the screen center (where the cursor was a white dot) onto the color wheel (where the cursor changed to a white rectangular box), and by clicking a button to confirm their choice. Once the selection was confirmed, both the color wheel and the response remained on the screen until the end of 4s. The color wheel was rotated by random degrees in each trial, thus avoiding confounding motor preparation with the reported color. Subjects were required to fixate throughout the trial.

The duration of one trial was either 18 s or 20 s, including an inter-trial interval (ITI) of 2 or 4 s (on average ITI = 3 s). A run was comprised of 50 trials in random order, with each of the 50 sample stimuli presented once as the cued stimulus and the not cued stimulus (fully randomized from each other). Three fMRI scanning sessions resulted in altogether 16 runs and 800 trials for the delayed estimation task per subject. Before the first scanning session, subjects were trained for half an hour with feedback on their responses.

### Perceptual Task

In the undelayed estimation task (‘perceptual task’), subjects reported a seen color on a concurrently presented color wheel. A trial began with the presentation of a sample stimulus in the middle of the screen (see **Figure 1b**). 500 ms after sample onset, the color wheel started to fade into view within a period of 350 ms. The color wheel was faded-in to minimize interference with the subjects’ perception of the color sample. The next trial started after an inter-trial interval of 2 or 4 s (on average ITI = 3 s). A trial was thus either 6 s or 8 s (on average 7 s), and a run consisted of 50 or 100 trials. Altogether 700 to 800 trials were conducted for the undelayed estimation task per subject.

### Category Naming and Identification Tasks

A pair of behavioral categorical tasks, color naming and identification, were performed in order to delineate the properties of color categories in our sample. The tasks were conducted in a dark behavioral lab using a keyboard and a mouse, after the completion of all fMRI sessions. Subjects had no time pressure as the next trial only started after they completed the current trial.

In the color naming task (**Figure 2b**), a list of seven common color names including ‘blue’, ‘pink’, ‘green’, ‘purple’, ‘orange’, ‘yellow’ and ‘red’ was shown next to a sample color. These chromatic color terms were selected based on Berlin and Kay’s eight basic color categories^29^ but ‘brown’ was excluded (see prior work^20^). Subjects were asked to select the term that best described the color stimulus by pressing the up or down button on the keyboard, and to confirm their choice by pressing enter. The order of the terms as well as the initial position of the cursor were randomized in each trial to minimize position bias. Six subjects completed 12 trials for each of the 50 color stimuli, while four subjects evaluated each stimulus 9 or 6 times (due to time constraints of the behavioral session).

In the color identification task (**Figure 2a**), subjects were required to mark the color wheel to identify the prototypical exemplar for of each of the seven color terms (see above). By pressing the left button of the mouse, they could confirm the color selection. The color wheel was rotated by random degrees in each trial to prevent association between the position and the color. Six subjects completed 90 identification trials for each category term, while four subjects evaluated each term 60 times.

### Stimuli

We used a set of 50 color samples taken from a circular color space with constant lightness (CIE LAB; center: a* = 0, b* = 0; radius = 38; L*=70; **Figure 1c**). Using a large number of different colors allowed us to finely sample variations in neural coding for stimuli in this circular space. A spectroradiometer (JETI spectraval 1501) was employed to measure L*a*b* values of each of the 50 generated colors, and to calibrate these parameters on different screens (the MRI monitor for the MRI session and the computer screen for the behavioral session). More specifically, we first calibrated the background gray color (used as the reference white point) to approximate a XYZ ratio of 1:1:1. Then, each color stimulus was measured and changed in multiple iterations to minimize the discrepancy to the chosen L*a*b* values.

### Data Acquisition

MRI data were acquired on a 12-channel Siemens 3 Tesla TIM-Trio scanner at the Berlin Center for Advanced Neuroimaging (BCAN). At the beginning of each scanning session, a high-resolution T1-weighted magnetization-prepared rapid gradient echo (MPRAGE) anatomical volume was collected (192 sagittal slices; repetition time TR = 1900 ms; echo time TE = 2.52 ms; flip angle = 9°; FOV = 256 mm). For acquisition of functional BOLD imaging, T2*-weighted echo planar images (EPI; 32 contiguous slices; TR = 2 s; TE = 30 ms; voxel size = 3×3×3 mm; matrix size = 64 × 64 × 32; slice gap = 0.6 mm; descending order; flip angle = 90°; FOV = 192 mm) were recorded covering the whole neocortex. Every trial was time-locked to the start of an EPI acquisition. For the memory task, 478 EPI scans were collected per run, and altogether 7648 scans were acquired over 16 runs per subject. For the perceptual task, data was acquired either in single (50 trials, 175 scans) or double (100 trials, 350 scans) runs. Overall, 2450 to 2800 functional scans were recorded per subject.

### Behavioral Analyses

To quantify the precision of color recall, we calculated the absolute error across trials separately for each condition (memory task and perceptual task) expressed in degrees of the 360° color-wheel. We also wanted to know whether subjects color reproductions were more biased during working memory as compared to a perceptual task (see ^20^). For this, bias in color reports was quantified by averaging recall errors for all repetitions of a given color (14-16 repetitions) and then averaging the absolute value of this individual bias across all colors. To control for the effect of random (color independent) recall errors on this bias measure, we permuted the color labels for each trial 1000 times and compared the resulting bias estimates against the biases computed from the real labels.

Data from the naming and the identification task was used to identify boundaries and prototypical colors for common color categories (for illustrative purposes, see **Figure 2d**). Naming data was minimally smoothed (Gaussian smoothing window, size 2, σ = 0.5) to minimize noise when determining boundaries between categories.

### Anatomical Regions of Interest

We focused our analyses on BOLD data from three regions of interest (ROIs) in visual cortex: V1, V4, and VO1. These ROIs (**Figure 3a**) were delineated based on high-resolution anatomical probabilistic maps^24^. These high-resolution probability maps were processed to obtain binary maps for every individual subject. First, the maps were deformed into the brain space of individual subjects using (inverse) normalization parameters obtained using unified segmentation. Then, the maps on the left and right hemispheres were collapsed and dorsal and ventral components were combined. We applied a mutual exclusion rule for all available probabilistic maps (see ^24^), such that every voxel could only be part of one ROI by selecting the ROI label with the highest probability. Finally, we threshold the resulting subject-level maps to exclude voxels with a probability lower than 10% to obtain a binary ROI map.

### fMRI Preprocessing

All fMRI analysis was conducted using SPM12^30^, cvManova^25^, and Matlab 2014b (Mathworks, Natick, MA). The acquired images were first converted from DICOM format to a SPM compatible format of NIfTI. Next, all functional images belonging to one subject were realigned and resliced to correct for head movement within and between runs. Then, the anatomical image was coregistered to the first functional image and subjected to unified segmentation (for inverse normalization).

### Sensory and Categorical Encoding Models

To estimate color-selectivity from spatially scattered and distinct response patterns of a population of voxels, we use two distinct encoding models: one sensory continuous and one categorical model (see **Figure 2c, e**). Encoding models capture the pattern of selectivity a voxel can be characterized as the weighted sum of a set of color-selective channels analogous to a neuron’s tuning curve. The *sensory* encoding model was characterized by six half-wave rectified cosine functions evenly distributed over the circular color space and raised to the power of six (**Error! Reference source not found.**). Such encoding models were used in prior work to model the selective neural response to orientation, spatial location and color^22,23,31,32^ and are intended to approximate single-unit tuning functions of *sensory* cortical neurons ^22,23^.

To model *categorical* neural representation, we developed a novel type of basis function of color (**Error! Reference source not found.c**) using empirical color categorization data (**Error! Reference source not found.ab**).

Six categorical basis functions captured the boundaries and prototypical colors of ‘blue’, ‘pink’, ‘green’, ‘purple’, ‘orange’, and ‘yellow’. To demarcate the boundaries of selectivity categories, we used the category naming data, and the category identification data was used to identify the prototypical exemplars of each category. The two corresponding probability distributions combining data from all subjects (**Figure 2c**, left) were normalized, so that the sum probability of each category equaled one. Then, we averaged the two probability distributions to create an encoding model capturing both the boundaries and the prototypical exemplars of each category. The resulting basis functions were normalized by dividing them by the highest value among all six channels.

Notably, this categorical encoding model does not require category-selective neuronal populations to exhibit an all-or-none response to any given stimulus. This is motivated by the behavioral data indicating that color categorization is probabilistic with the same hue being assigned different color categories in different trials in the naming task. Further, we anticipated that categorically color-selective voxels respond strongest to prototypical exemplars of a given color category. Finally, fitting the six basis functions simultaneously allows any voxel to have positive weights for multiple color categories as it might contain neurons selective for multiple categories.

Thus, we created two distinct encoding models: (1) A *sensory* model used in prior work to resemble the tuning of sensory neurons while carrying no information about the delineations of common color categories, and (2) a categorical encoding model informed by empirical categorization data.

### Multivariate Pattern Analysis

To test which of these models best explained mnemonic activity patterns during the working memory delay, we combined these two encoding models with a recently-developed form of multivariate pattern analysis (MVPA), cross-validated multivariate analysis of variance (MANOVA^25^). The analysis was performed on a set of selected voxels within three regions of interest (ROIs): V1, V4, VO1 (see **Figure 3a**). For this, we first estimated parameters (i.e., betas) for multivariate generalized linear models (MGLM) separately for each condition (memory and perceptual) and encoding model (sensory and categorical) which modelled sample colors as a set of six parametric modulations representing the six basis functions per model.

For the memory task, we used five finite impulse response (FIR) regressors to represent the 10 s delay-period (5 fMRI scans at a TR of 2 s). The design matrix modelled the 478 scans per run using 36 regressors (7 stimulus-based regressors [1 constant and 6 basis functions] x 5 FIR bins + 1 run-wise constant). Separate design matrices and MGLMs used basis functions from the *sensory* and the *categorical* encoding model.

For the perceptual task, but the 4 s stimulus presentation was represented by a canonical hemodynamic response function (HRF, duration = 4 s), which was time-locked to the stimulus presentation’s onset. The design matrix captured each run (either 175 or 350 scans) using 8 regressors (7 stimulus-based regressors [1 constant and 6 basis functions] x 1 HRF + 1 run-wise constant). Again, separate design matrices and MGLMs used basis functions from the *sensory* and the *categorical* encoding model. Parameter estimates for all models were estimated using standard SPM parameters, but parametric modulations were not orthogonalized and serial correlations corrections were omitted. Next, we estimated the variance explained by these models by contrasting each neighboring pair of basis functions (BF 1 vs BF 2; BF 2 vs BF3; BF 3 vs BF 4...) separately for each time point (for the memory task). For the memory task, the overall contrast matrix for a given run was comprised of 35 columns representing 35 regressors (six BF-based and one stimulus-based regressors, each in five FIRs bins) and 25 rows representing 25 contrasts (five contrasts between six BF-based regressors, each in five FIR bins). For the perceptual task, the contrast matrix for a given run had 7 columns representing 7 regressors (six BF-based and one stimulus-based regressors, each in one HRF bin) and 5 rows representing 5 contrasts (five contrasts between six BF-based regressors). The null hypothesis, here, is that in a given set of voxels there are no differences in the parameter estimates for the six basis functions in a given model. Rejecting this null indicates that this subset of data carries information about sample color in a given trial. The resulting pattern distinctness D (see ^25,33,34^) reflects the variance of the neural data explained by the respective model. To assess statistical significance against chance-level (D = 0) and to compare models against each other as well as model-by-task interactions, we used a nonparametric bootstrapping testing group effects by random resampling 10e^5^ times ^35–37^. We corrected the resulting confidence intervals for the multiple comparisons in the three different ROIs using Bonferroni correction (resulting in an effective confidence interval of 98.33 %).

## Data Availability

All data analyzed in the current study are available from the corresponding author upon request.

